# Characteristics of ectopic alveolar basal cells relative to airway basal cells in fibrosis

**DOI:** 10.1101/2024.09.24.614660

**Authors:** Sabrina Blumer, Petra Khan, Julien Roux, Nataliia Artysh, Linda Plappert, Antje Prasse, Katrin E. Hostettler

## Abstract

**Rationale:** Basal cells (BC) appear ectopically within the lung parenchyma of interstitial lung disease (ILD) patients, potentially through migration of airway BC or though trans-differentiation of alveolar epithelial type 2 (AT2) cells. The exact origin and function of these ectopic alveolar BC remains elusive. By comparing ectopic alveolar to “classical” airway BC, we aimed to get a better understanding of the origin and characteristics of alveolar BC in ILD.

**Methods:** Alveolar and airway BC were isolated from transbronchial and airway mucosal biopsies, respectively, from the same ILD patients and expanded in culture. Samples were analyzed by single cell RNA sequencing (scRNA-seq), TaqMan RT-PCR, and immunochemistry.

**Results:** scRNA-seq analysis revealed several differences in gene expression that suggested a shift to a more mesenchymal-like phenotype and a decrease in keratinization genes in alveolar compared to airway BC. Specific AT2 cell marker genes were not expressed in either BC type. While the morphology, wound repair and proliferation capacities of BC from both origins were not significantly different, alveolar BC formed significantly fewer organoids, expressing more MUC5B. After instillation into bleomycin-injured mice, alveolar and airway BC showed similar engraftment, differentiation capacity and effects on fibrosis.

**Conclusion:** Despite similar overall functionality in vitro and after instillation into bleomycin-injured mice, alveolar and airway BC differed in their transcriptomes and in their capacities to form and to differentiate in organoids. Our data provide no evidence to support their potential derivation from AT2 cells.

**Take home message:** Alveolar and airway basal cells differ in their transcriptomes and in their capacities to form and to differentiate in organoids, although with no indication of an AT2 cell origin.

## Introduction

In interstitial lung diseases (ILD) basal cells (BC) accumulate within the alveolar space, where they replace resident alveolar epithelial cells and contribute to pathological honeycomb cysts (HC) formation [1–3]. In the healthy lung, alveoli are lined with alveolar epithelial type (AT)1 and 2 cells, whereas BC are restricted to the airways. AT2 cells are critical for alveolar homeostasis through pulmonary surfactant production and their ability to self-renew and differentiate into AT1 cells [4]. BC maintain airway homeostasis by self-renewal and the differentiation to other airway epithelial cells such as ciliated, or secretory epithelial cells [5].

The origin of ectopic BC in the alveolar space in ILD remains elusive. BC are present throughout the airways all the way to the terminal bronchioles in human [6]. It is therefore possible that alveolar BC derive from migrated airway BC. Alternatively, studies showed the possibility for AT2 cells to trans-differentiate into BC *in vitro* [7, 8], constituting an alternative explanation for the origin of the alveolar BC pool in ILD.

Some studies suggested a beneficial role of the ectopic BC in ILD in regenerating the injured alveolar epithelium through trans-differentiation to AT2 cells in mice [9, 10]. However, other studies did not find evidence for a contribution of ectopic BC to the AT2 cell pool [11, 12], and suggested that ectopic BC could even play a detrimental role because their abundance correlated with increased mortality and HC formation in ILD [3, 13].

By comparing alveolar BC to their airway counterpart after isolating these two respective BC types from ILD patients and expanding them in culture, we aimed to get a better understanding of the origin and disease-specific characteristics of ectopic BC, which may help to further unravel the pathomechanism of fibrotic ILD.

## Materials and Methods

### Basal cell culture

Alveolar BC were cultured from transbronchial biopsies (TBB) and airway BC from airway mucosal biopsies (MB) [14–16]. Briefly, TBB or airway MB derived from the same ILD patient were placed into plastic dishes containing an epithelial cell-specific growth medium (Cnt-PR-A). After 5-7 days, tissue was removed and the outgrown cells expanded. Experiments were performed with BC from tissue of several different ILD patients as indicated by the n-number. The local ethical committee of the University Hospital, Basel, Switzerland (EKBB05/06) approved the use of human tissue sections and the culture of human primary lung cells. ILD patient characteristics and material used in this study are outlined in supplement table 1 and 2.

### TaqMan RT-PCR, Immunochemistry

TaqMan RT-PCR, immunocytochemistry (ICC)/immunofluorescence (IF), immunohistochemistry (IHC)/IF or hematoxylin and eosin (H&E) stainings were performed as previously described [15–17]. Primers and antibodies are listed in supplement table 2.

### Single-cell RNA-sequencing

scRNA-seq was performed on cultured airway and alveolar BC from three different ILD patients (Patient-ID 01, 02, and 03) using the 10X Genomics technology [17]. Detailed scRNA-seq analysis methods can be viewed in supplement document 1.

### Air liquid interface (ALI), organoids, proliferation and epithelial wound repair

BC ALI- and organoid culture as well as proliferation and epithelial wound repair assays were performed as previously [16].

### Intratracheal administration of human BC into bleomycin-challenged mice

The Hannover Medical School, Germany performed all mouse procedures in accordance with the German law for animal protection and the European Directive 2010/63/EU and an approval by the Lower Saxony State Office for Consumer Protection and Food Safety in Oldenburg/Germany (LAVES); AZ: 33.12-42502-04-15/1896 and AZ: 33.19-42502-04-15/2017), AZ:33.12-42502-04-17/2612). Human alveolar and airway BC were transduced with a lentiviral vector for firefly-luciferase and enhanced green fluorescence protein (eGFP) [13, 16]. NRG mice (n=9) received bleomycin (1.2 mg/kg) intratracheally. After three days, three mice were intratracheally injected with human alveolar BC, and three mice with human airway BC (both at 0.3 X 10^5^ cells per mouse). Firefly-luciferase activity was measured as previously described [13, 16]. At day 21, mice were sacrificed and mouse lungs were formalin-fixed and paraffin embedded (FFPE). Fibrosis scoring was performed on lung tissue of mice treated with bleomycin only (n=3), bleomycin + human alveolar (n=3), or + human airway (n=3) BC [18]: 20 non-overlapping 10x fields per sample were scored as followed: 0L=Lno fibrosis, 1L=Lmild thickening of alveolar septa, 2L=L<10%-, 3L=L10–20%, 4L=L20–40%-, 5L=L40–60%-, 6L=L60–80%, 7L=L>80% fibrotic area or 8L=Lcomplete fibrosis. The scores of all fields were then averaged for each sample.

### Statistics

Paired or unpaired t-tests were performed using GraphPad Prism software version 9.1.1. p values < 0.05 were considered as statistically significant. Data were presented as mean ± SEM.

## Results

### BC characteristics

In airway MB, KRT5+/p63+ BC were lining the basement membrane of the airway epithelium, whereas in TBB, they were found scattered within the lung parenchyma (Figure 1 a). Alveolar and airway BC, isolated and cultured as illustrated (Figure 1 b), were indistinguishable by their morphology (Figure 1 c) and expressed similar levels of BC markers (KRT5, KRT14, KRT17, TP63) both at the RNA and protein level (Figure 1 d). Mesenchymal (CDH2)- or secretory epithelial (SCGB1A1) cell markers were absent or expressed at very low levels (Figure 1 d).

**Figure 1:**
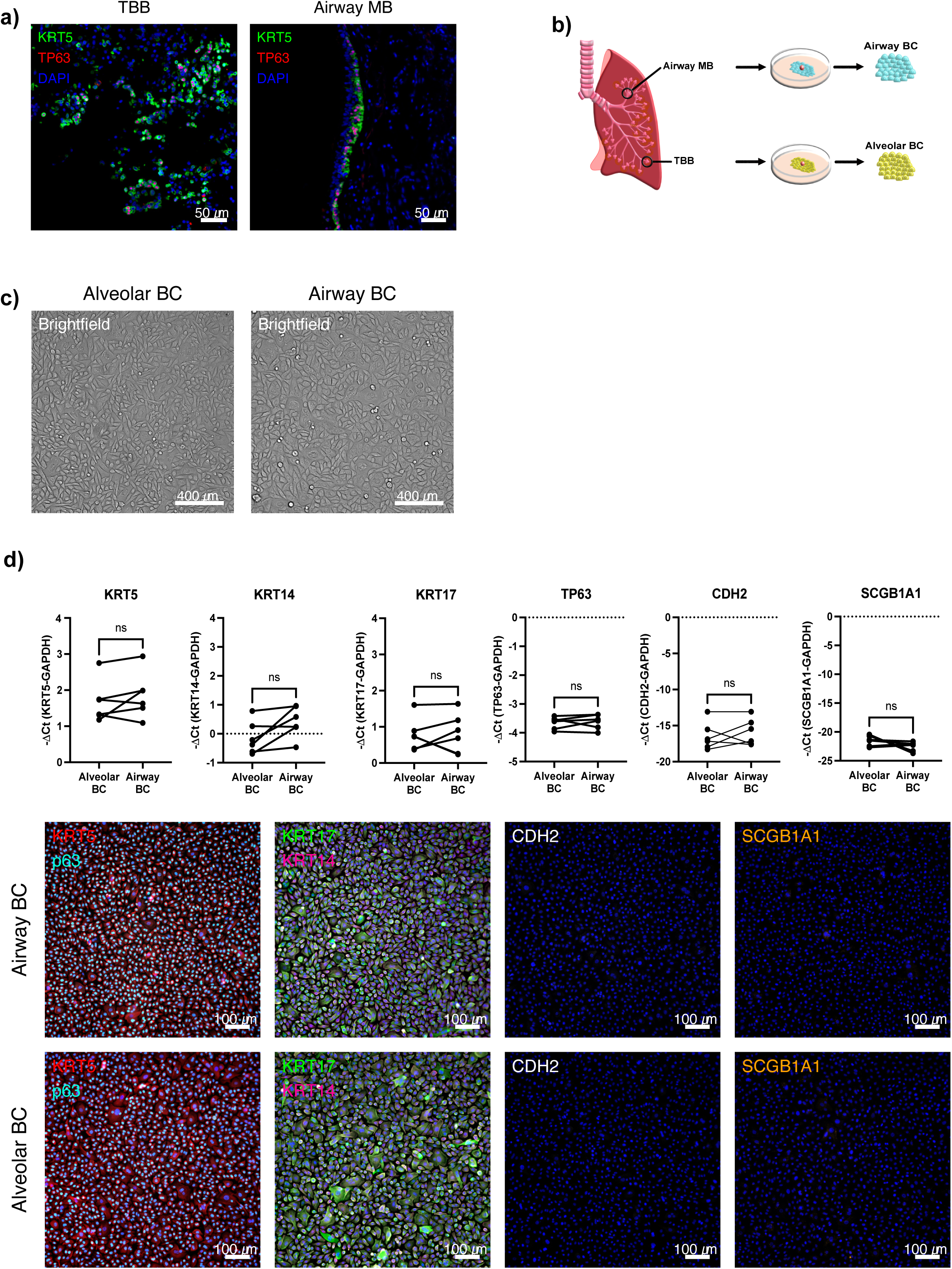
Cultured alveolar and airway BC do not differ in their morphology or BC marker expression. KRT5+ and TP63+ BC in formalin fixed paraffin embedded transbronchial biopsies or airway mucosal biopsies of the same ILD patient (n=5) (a). To culture alveolar or airway BC, tissue specimens were placed in cell culture dishes containing Cnt-PR-A as illustrated in b. Representative phase contrast images showing the morphology of confluent alveolar or airway BC (n=5) (c). RNA expression of KRT5, KRT14, KRT17, TP63, CDH2 or SCGB1A1 (n=6; paired t-test, ns indicates not significant) and representative IF stainings for KRT5/p63, KRT17/KRT14, CDH2, SCGB1A1 KRT5/TP63 (n=3) (d) in cultured alveolar and airway BC.

### BC transcriptome

scRNA-seq was performed on cultured alveolar and airway BC from 3 ILD patients. After quality control and filtering of cells, 34,435 cells were retained (4,500 to 6,975 cells per sample). The major source of variation across single cells was the individual origin of the sample (Donors 1-3), followed by the origin of the BC samples (TBB or airway MB) (Figure 2 a; Supplement figure 1). After removal of these systematic sources of variation, the cells were grouped into 7 different clusters encompassing between 7.6% and 20% of the cells (Figure 2 b). Each cluster was annotated by comparing our cells to published scRNA-seq atlases of the lung (Figure 2 c [19]; Supplement Figure 2 [1, 2, 7, 20–25]): the majority of cells in clusters 2-6 (84 to 97%) showed closest transcriptomic similarities to the BC subset 1 (canonical BC) described in [19]. Most cells (86%) from cluster 7 closely matched to BC subset 2 (cycling BC) [19]. Last, a large fraction (39%) of cells from cluster 1 matched to BC subset 3 (secretory-primed) [19]. This annotation was supported by inspection of the expression of the best markers across clusters (Figure 2 d) and known marker genes (Supplement Figure 3). More specifically, cluster 7 displayed high expression of proliferation genes such as *PTTG1*, *BIRC5*, *HMGB2*, *MIK67*, *CENPF* and *TOP2A* (Figure 2 d). Secretory-primed BC from cluster 1 were enriched with transcripts of members of the serpin family, *KRT16*, or *CSTA* (Figure 2 d). The markers of the other clusters seemed to illustrate a differentiation gradient from cluster 6 to the secretory-primed cluster 1, analogous to previously reported [15]. In particular cells from cluster 6 were enriched with transcripts highly expressed in undifferentiated BC [26] such as *AREG*, *CAV1*, *THBS1*, or *ITGB4*, whereas transcripts indicating BC to secretory epithelial cell differentiation (*CSTA*, *KRT16*, *SERPINB2*) [27] were expressed at low levels. Their expression increased in cluster 3 and 4 and was highest in cluster 1 (Figure 2 d). Despite expressing lower levels of *AREG*, *CAV1*, *THBS1* and *ITGB4* than cluster 6, cluster 2 showed also lower levels of *CSTA*, *KRT16*, *SERPINB2* than cluster 3 and 4. Instead, high levels of expression of several metallothioneins (e.g., *MT1X*, *MT2A*, *MT1E*), *FOS*, *KRT15*, or *AGR2* were observed, which were in part shared with cluster 3 (Figure 2 d). Cluster 5 showed high expression of the mesenchymal markers *FN1* and *VIM* (Figure 2 d), but it was predominantly composed of cells from donor 1 (74% of cells, Supplement figure 1 a). Of note, there was overall no or very low expression of specific AT2 cell markers (e.g., *SFTPA-D*, *ABCA3*) across cells in our dataset (not shown).

**Figure 2:**
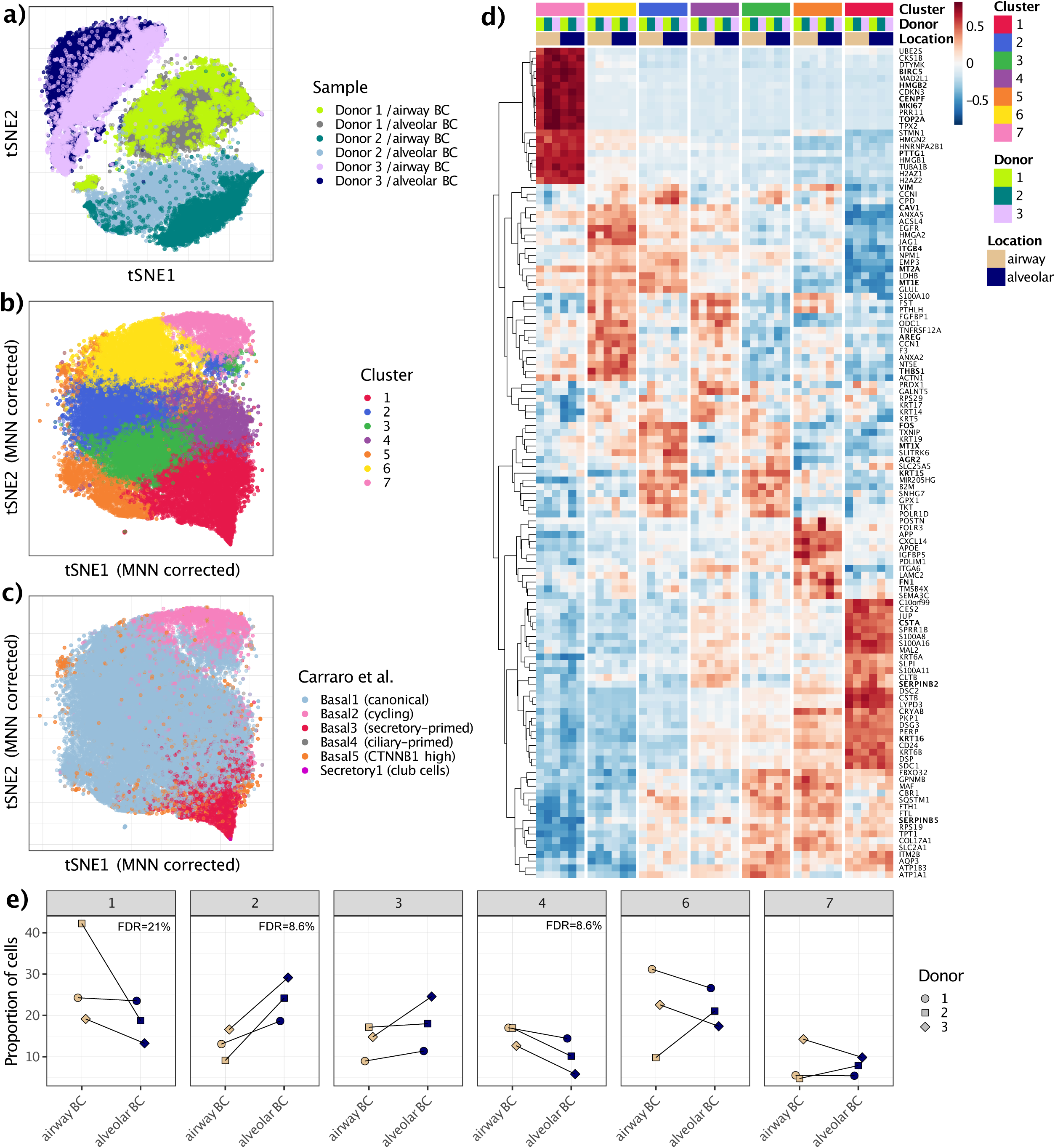
Alveolar and airway BC differ in their transcriptome. T-SNE plot (uncorrected data) showing cultured alveolar and airway BC derived from donor 1-3 (a). T-SNE (after MNN correction) showing the separation of cells into 7 clusters (b). T-SNE showing the best-matching reference label in the transcriptome comparison to a publicly available scRNA-seq dataset [41] (c). Heatmap of the relative expression across clusters, donors, and sample location of the strongest cluster-specific genes. The log- normalized expression levels were averaged across cells from the same cluster, donor, and sample location, corrected for donor-specific effects to mirror the MNN correction performed before clustering and the inclusion of donor as a covariate in the differential expression analysis, and averaged values were then centered and scaled per gene. Marker genes are highlighted in bold (d). Proportion of alveolar and airway BC across donors in clusters 1, 2, 3, 4, 6, and 7 (e).

Comparing the relative abundance of alveolar compared to airway BC across clusters (excluding cluster 5 including mostly cells from donor 1), we observed a significantly higher abundance of alveolar BC in cluster 2 (FDR=8.6%), along with a reduced abundance of alveolar BC in cluster 4 (FDR=8.6%) and cluster 1 (FDR=21%) (Figure 2 e). These results suggest an enrichment for an altered differentiation trajectory among alveolar BC (cluster 6 to clusters 2 and 3) relative to the “classical” differentiation trajectory (cluster 6 to cluster 4 and then to cluster 1) enriched among airway BC.

Next, we tested for differences in expression state of alveolar compared to airway BC. Testing within each cluster separately allowed us to account for the above-described differences in abundance, while the paired design allowed us to account for donor-specific effects. At an FDR of 1%, there were in total 40 genes significantly up or down-regulated in alveolar compared to airway BC in at least one of the clusters (Figure 3 a). Cluster 2 and 6 yielded the highest amount of differentially expressed genes (Figure 3 b; differential expression in other clusters is shown in Supplement Figure 4). Genes involved in response to hypoxia (*PLIN2*, *S100A4*), KRAS activation/mutation (*LMO3*, *IL33*), ECM organization (*SPOCK3*, *MXRA7*), as well as genes involved in glycolysis (*EGLN3*) were up-regulated in alveolar BC, whereas genes involved in interferon response (*IFI27*, *HLA-DRB1*, *BST2*) were down-regulated. Interestingly, although not significant, the BC marker genes *KRT5*, *KRT14, KRT17* and *TP63* were all down-regulated (Supplement Figure 3), whereas *KRT15* and *KRT19* were up-regulated, along with mesenchymal markers such as *FN1*, *COL1A1* or *VIM*.

**Figure 3:**
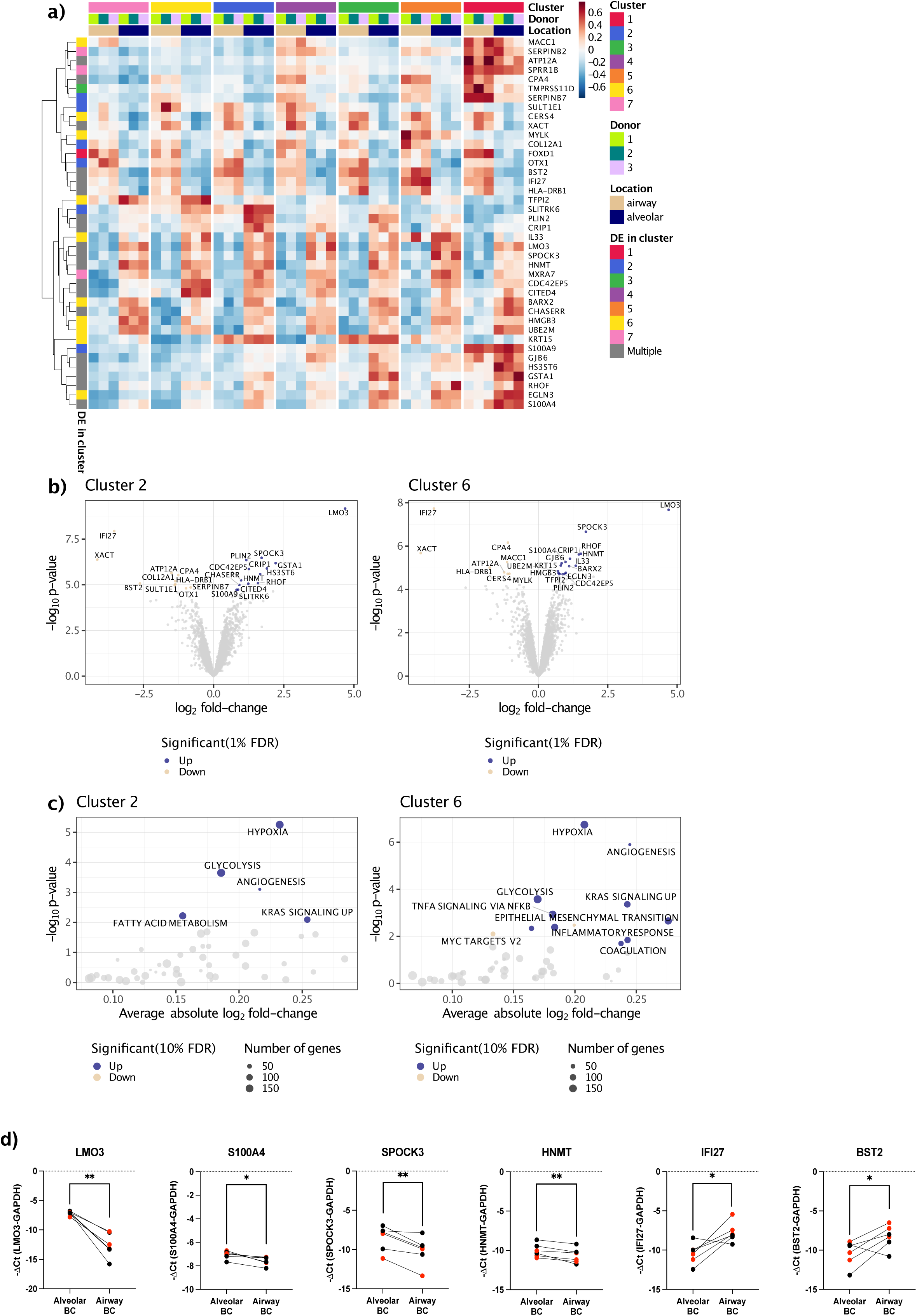
Differential genes and pathways in alveolar versus airway BC. Heatmap showing the differential expressed genes in alveolar versus airway BC across all tested clusters. The plotted relative expression values were calculated similarly to Figure 2D. Annotation colors for rows indicate in which specific cluster each gene was found significantly differentially expressed at an FDR threshold of 1% (a). Volcano plots showing the differential expression analysis results between alveolar and airway BC in cluster 2 and cluster 6 (b). Genes colored (beige=decreased, and blue=increased in alveolar BC relative to airway BC) are significant at an FDR threshold of 1% (b). Functional enrichment analysis amongst differentially expressed genes in cluster 2 and cluster 6 (c) using MSigDB Hallmark gene sets. Six differentially expressed genes (*LMO3*, *S100A4*, *SPOCK3*, *HNMT*, *IFI27*, *BST2*) were selected and analyzed by Taqman RT-qPCR (d). Data points in red represent results from cells derived from donor 1-3. Data points in black were derived from cells of three additional donors (paired t-test, ** indicates p < 0.01, * indicates p < 0.05).

Functional enrichment analysis using the MSigDB Hallmark pathways (Figure 3 c for clusters 2 and 6; Supplement Figure 5 for clusters 1, 3, 4 and 7) showed that pathways related to hypoxia, glycolysis, angiogenesis, KRAS signaling, and epithelial to mesenchymal transition (EMT) were up-regulated in alveolar BC. Furthermore, Reactome pathway analysis revealed an up-regulation of pathways such as “ECM organization”, “collagen formation”, “collagen biosynthesis and modifying enzymes”, or “activation of matrix metalloproteinases” in alveolar BC (Supplement Figure 6). In line with the above-described trend of keratin genes, the pathway “Keratinization” was significantly down-regulated in alveolar BC in cluster 1, 6, and 7 (Supplement Figure 6).

The differential expression of selected genes was validated in BC from donor 1-3 and three additional donors by TaqMan RT-PCR, confirming the up-regulation of *LMO3*, *S100A4*, *SPOCK3*, *HNMT* and down-regulation of *IFI27*, *BST2* in alveolar relative to airway BC (Figure 3 d).

### BC proliferation and wound healing capacities

Alveolar and airway BC numbers significantly increased between 0-48 h (Figure 4 a) and closed a scratch after 6 hours (Figure 4 b) with no difference in cell growth rate (Figure 4 a) or wound healing capacity (Figure 4 b) between alveolar and airway BC.

**Figure 4:**
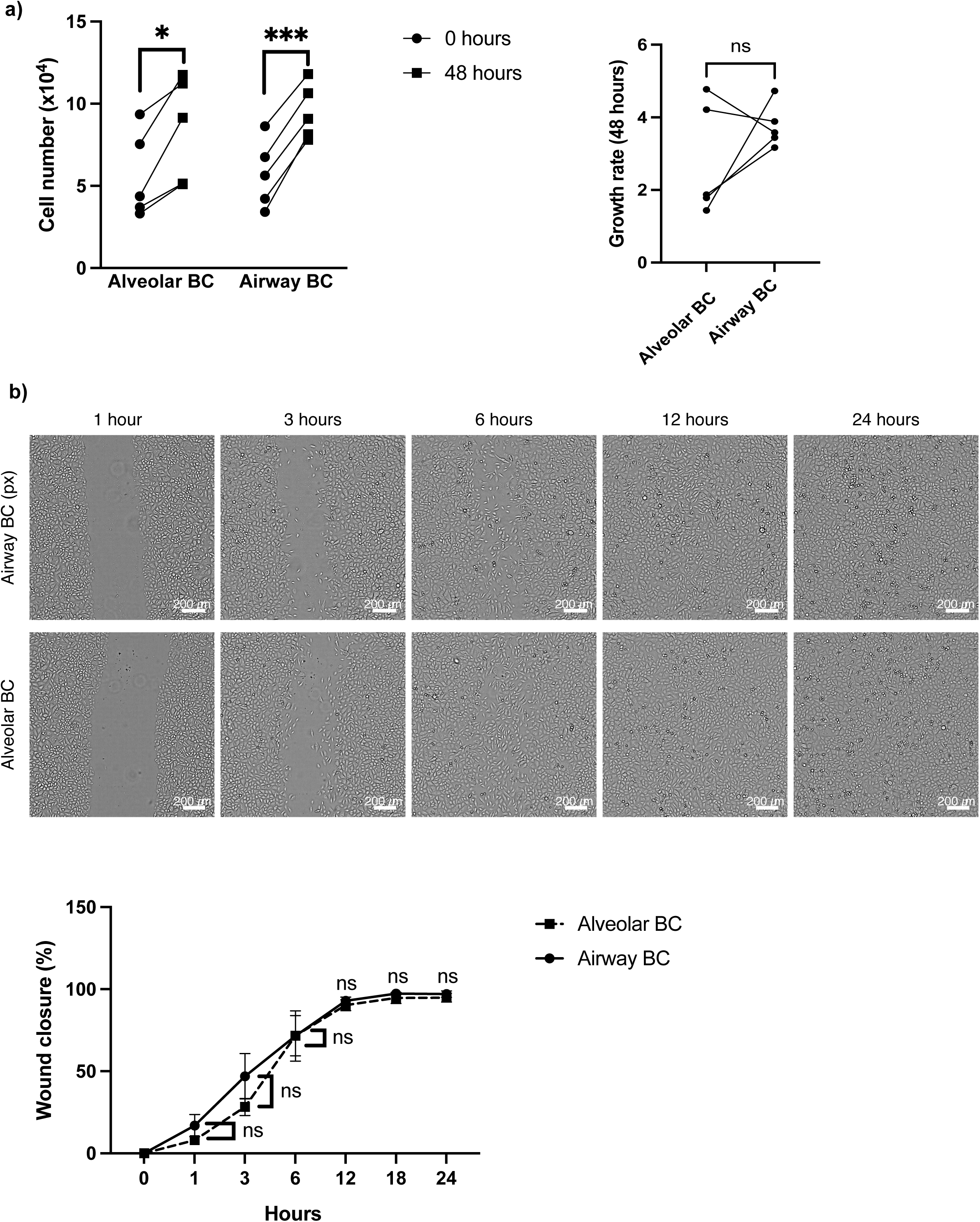
Alveolar and airway basal cells display similar proliferation and wound healing capacities. Automated cell counts of cultured alveolar and airway BC at 0 and 48 hours (paired t-test, *** indicates p < 0.001, * indicates p < 0.05) and their growth rate (cell number at day 48 - cell number at day 0) comparing alveolar to airway BC (n=4; paired t-test, ns indicates not significant) (a). Representative phase contrast images and graph displaying alveolar and airway BC wound closure (%) between 0-24 h (mean ± SEM, paired t-test ns indicates not significant) (b).

### BC differentiation capacities

Alveolar and airway BC cultured on an ALI (Figure 5 a) differentiated to secretory and ciliated epithelial cells, reflected by a significant increase of secretory (*SCGB1A1*, *MUC5AC*, *MUC5B)* and ciliated (*FOXJ1)* epithelial cell marker expression levels between day 0 and 23 (Figure 5 b). Marker expression levels did not significantly differ between alveolar and airway BC at day 0 or 23 (Figure 5 b). At the protein level, BC did not express AcTub (ciliated epithelial cell marker), MUC5AC, nor SCGB1A1 at day 0 (Figure 4 c), while after 23 days these markers were detected in alveolar and airway BC on PFA-fixed (Figure 5 c) and/or cryo-sectioned (Figure 5 d) ALI membranes.

**Figure 5:**
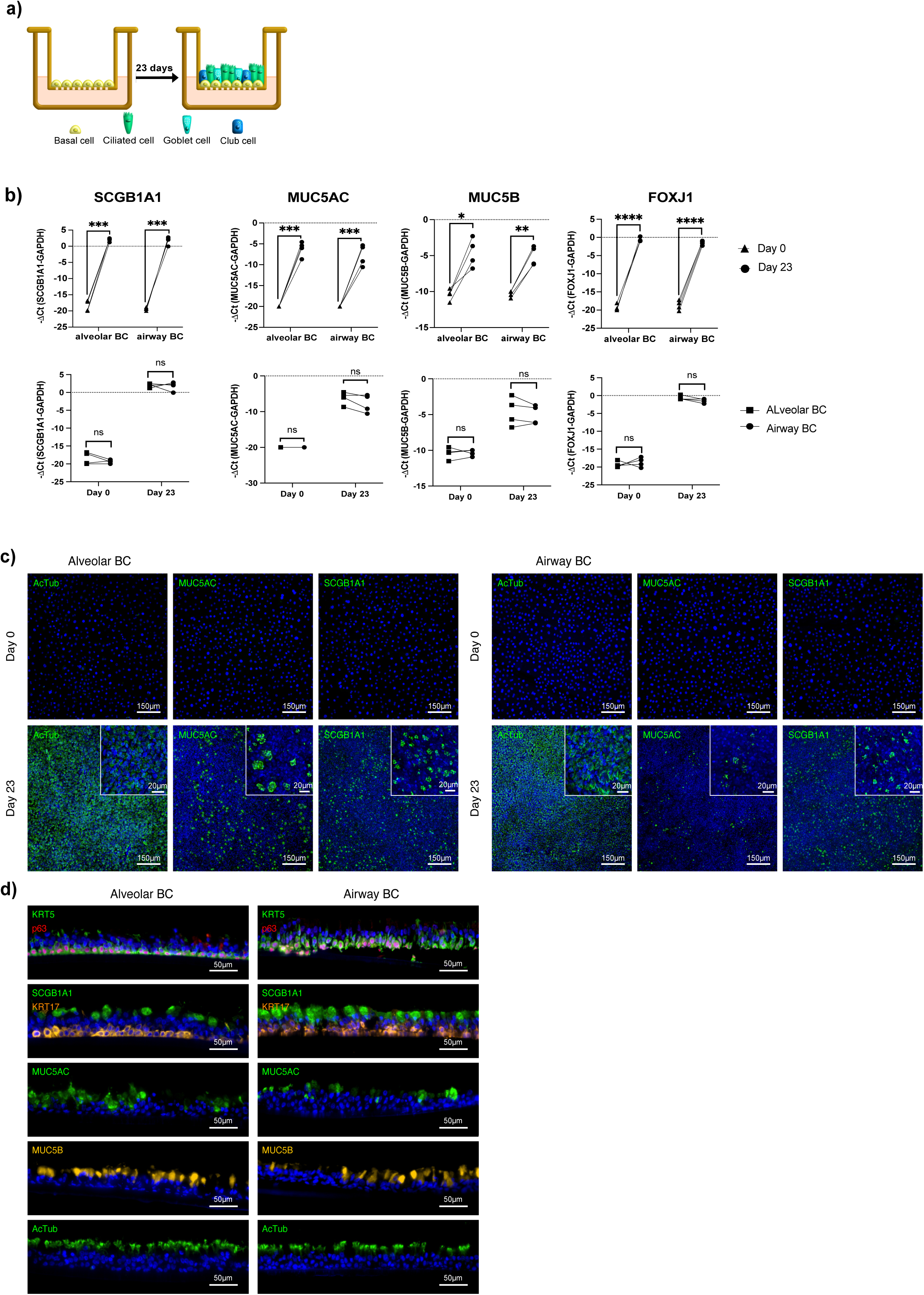
Alveolar and airway basal cells have similar differentiation capacities. Illustration of alveolar and airway BC ALI culture (a). RNA expression levels (-ΔCt) of KRT5, SCGB1A1, MUC5AC, MUC5B, or FOXJ1 in ALI-cultured alveolar and airway BC (n=4) at day 0 and 23 (paired t-test, **** indicates p < 0.0001, *** indicates p < 0.001, ** indicates p < 0.01, * indicates p < 0.05) and comparing their expression in alveolar versus airway BC (paired t-test, ns indicates not significant) (b). When the target genes were expressed at levels too low to be detected after 40 cycles, the missing -ΔCt values were arbitrarily set to −20 (b). Representative ICC/IF images of alveolar and airway BC (n=3) incubated with antibodies detecting acetylated-tubulin (AcTub), MUC5AC or SCGB1A1 at day 0 and 23 (c). Representative images of cryosections incubated with antibodies detecting KRT5, p63, SCGB1A1, KRT17, MUC5AC, MUC5B, or AcTub of ALI-cultured alveolar or airway BC (n=3) at day 23 (d).

BC cultured in Matrigel (Figure 6 a) formed organoids (>50µm diameter) after 7 days, which grew in size and number over a period of 21 days (Figure 6 b). After 20 days, alveolar BC formed between 45 and 138 organoids/mm^3^, significantly fewer than the 63 to 162 organoids/mm^3^ formed by airway BC (Figure 6 c). The size of alveolar (66-122 µm) and airway (73-105 µm) BC-derived organoids was not significantly different (Figure 6 c). After 21 days, cells in alveolar and airway BC-derived organoids expressed similar RNA levels of *TP63*, *KRT5*, *KRT17*, *SCGB1A1*, and *MUC5AC. MUC5B* levels were significantly higher in alveolar BC-derived organoids and FOXJ1 levels showed a non-significant upward trend (Figure 6 d). Marker protein levels were expressed in both alveolar and airway BC, with several cells still expressing basal cell markers (KRT5, p63, KRT17). Differentiated cells primarily expressed MUC5AC, and/or MUC5B, while SCGB1A1 or AcTub proteins were detected in only a view cells (Figure 6 d).

**Figure 6:**
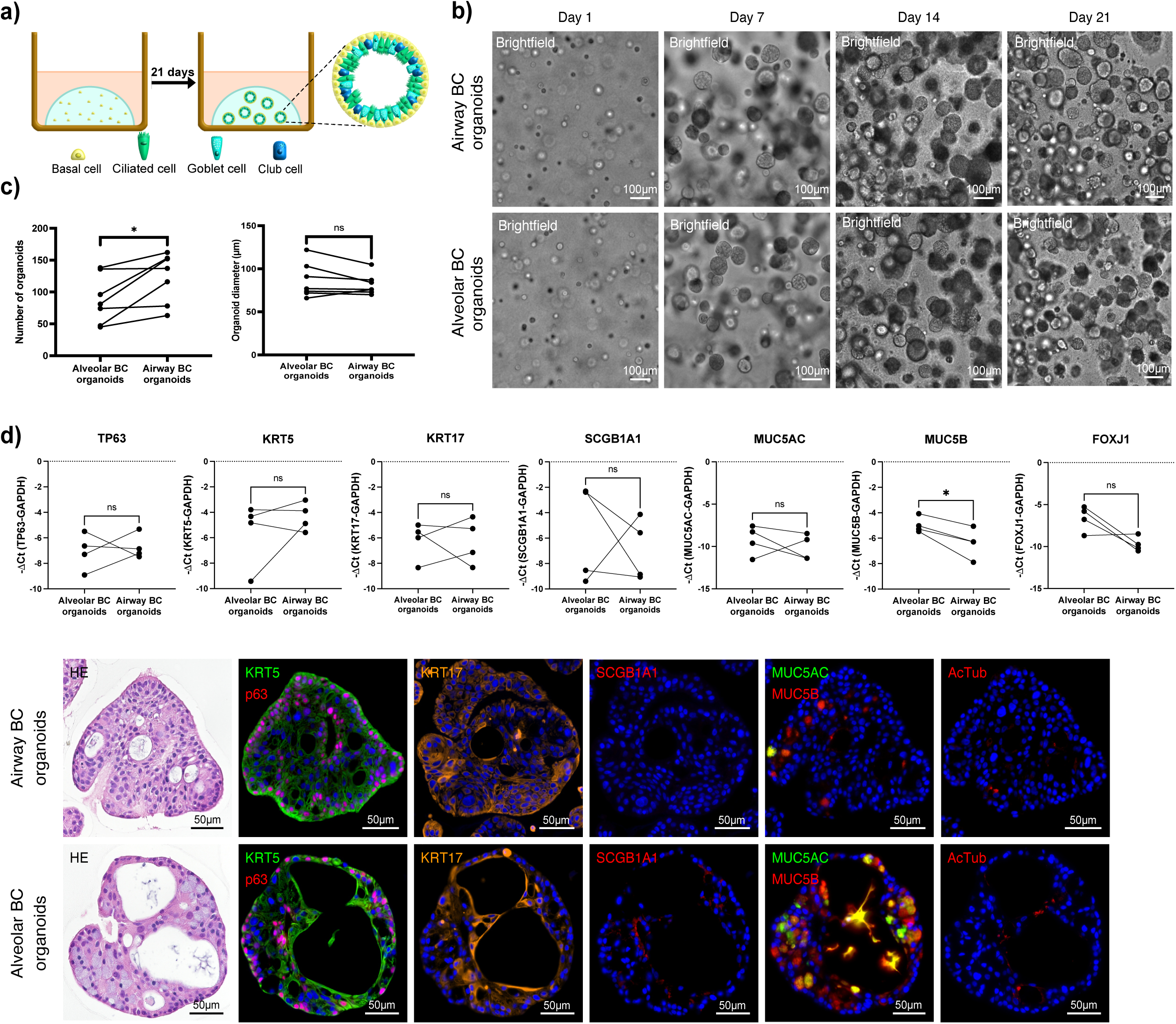
Alveolar and airway basal cells form organoids containing basal, secretory- and ciliated epithelial cells. Alveolar and airway BC were resuspended in Matrigel and cultured for 21 days as illustrated (a). Representative phase contrast images of alveolar or airway BC (n=7) organoid formation at day 2, 7, 14, and 21 (b). Number of alveolar and airway BC-derived organoids/mm^3^ and the distribution of their diameters (>50μm) after 20 days (n=7; paired t-test, * values ≤ 0.05, ns=not significant) (c). RNA expression levels (-ΔCt) of TP63, KRT5, KRT17, SCGB1A1, MUC5AC, MUC5B, or FOXJ1 in organoids after 21 days (n=4; paired t-test, * values≤ 0.05, ns=not significant) and representative H&E and ICC/IF images of alveolar or airway BC-derived organoids (n=3) incubated with antibodies detecting KRT17, KRT5, p63, SCGB1A1, AcTub, MUC5AC, or MUC5B (d).

### Human BC in bleomycin-challenged mice

Mice were treated as outlined in figure 7 a. BC engrafted and proliferated, which was determined by increasing luciferase expression (Figure 7 b). Bleomycin induced mild fibrotic changes in the mouse lungs, which were slightly increased when alveolar or airway BC were injected (Figure 7 c), but according to fibrosis scores the increase was not significant (Figure 7 c). Co-stainings with human-specific HNA and basal-(KRT5, KRT17), secretory-(MUC5AC, MUC5B, SCGB1A1), or ciliated (AcTub) epithelial cell markers, revealed that human BC maintained their BC identity or differentiated to MUC5AC+, MUC5B+, or SCGB1A1+ cells in mouse lungs (Figure 7 d). We did not observe differences in fibrosis scores or differentiation capacity within mouse lung tissue, which were injected with either human alveolar or airway BC (Figure 7 c, d).

**Figure 7:**
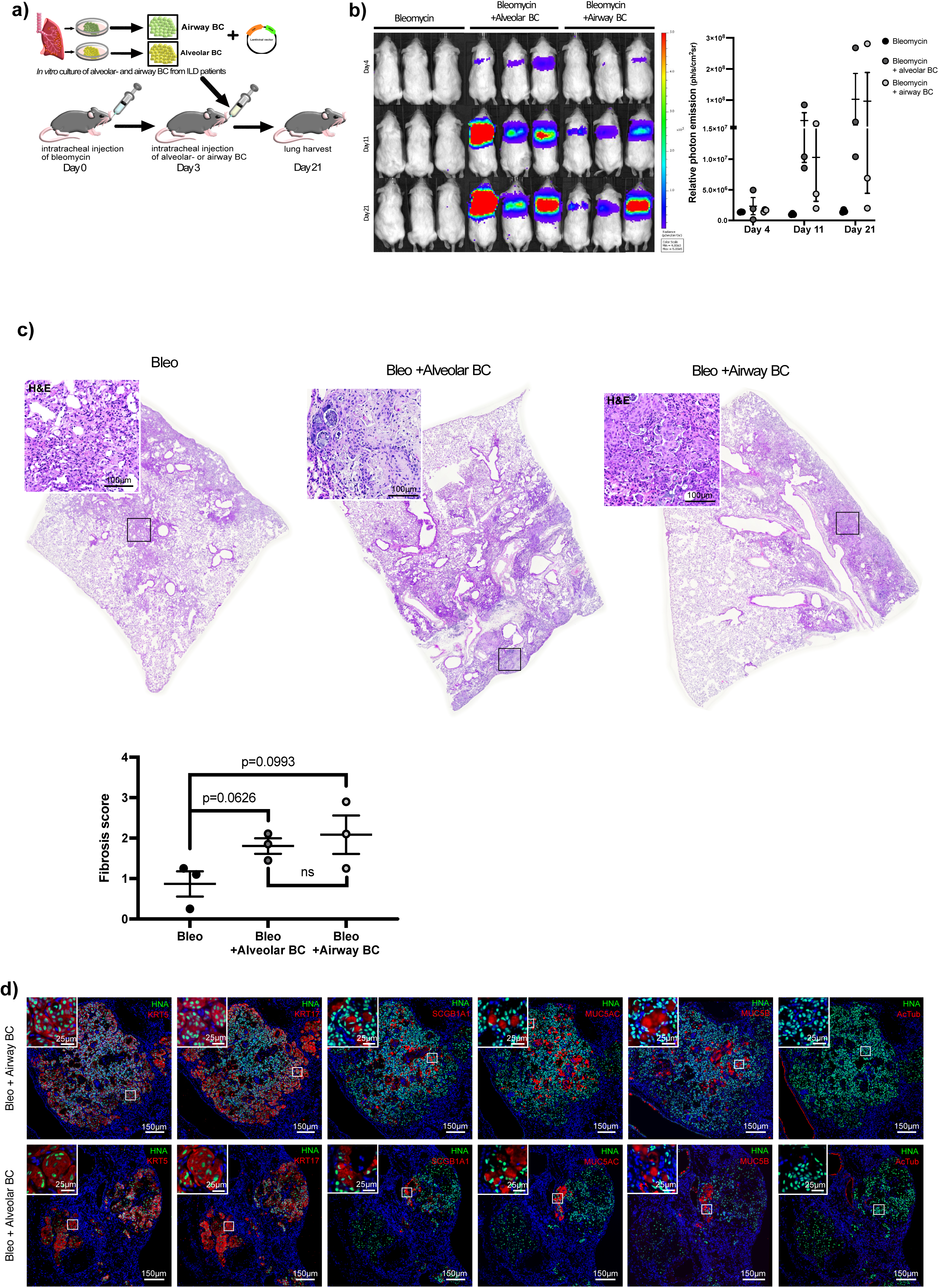
Human alveolar or airway basal cells in bleomycin-challenged mouse lungs. NRG mice were treated with bleomycin (n=3) +/- human alveolar (n=3) or airway (n=3) BC carrying a luciferase encoding vector as illustrated (a). Luciferase expression in mice treated with bleomycin +/- human alveolar or airway BC carrying a luciferase encoding vector and the corresponding bioluminescence measurements at day 4, 11, and 21 (b). Representative images of H&E stainings and fibrosis scores for mice treated with bleomycin +/- human alveolar or airway BC (unpaired t-test, Bleo versus Bleo+ alveolar or airway BC; paired t-test for Bleo+ alveolar BC versus Bleo+ airway BC) (c). Representative IHC/IF images of mouse lung tissue (mice treated with bleomycin + human alveolar or airway BC) incubated with antibodies detecting human-specific HNA with or without KRT5, KRT17, SCGB1A1, MUC5AC, MUC5B, or AcTub. Nuclei were counter-stained with DAPI. White squares indicate regions imaged at higher magnification (d).

## Discussion

To investigate the origin and characteristics of ectopic alveolar BC in ILD, we made use of a previously established *in vitro* BC culture method [16] allowing us to isolate and expand alveolar and airway BC from the same ILD patients in sufficient numbers. Our data show that BC from the different origins display similar morphology, proliferation and wound healing capacities *in vitro,* as well as comparable engraftment and differentiation abilities when instilled into bleomycin-challenged mice. Despite many similarities, cultured alveolar and airway BC significantly differed in their transcriptomes, and in their potential to form and to differentiate in organoids, although with no indication of an AT2 cell origin.

Transcriptomic differences between alveolar and airway BC included an up-regulation of genes associated with fibrosis-related pathways such as hypoxia, glycolysis, or KRAS signalling in alveolar BC. Hypoxia is increased in fibrotic areas of the ILD lung [28–30], and cellular glycolysis [31] and KRAS signalling [32] are activated under hypoxic conditions. Ectopic BC specifically accumulate in hypoxic regions of the ILD lung parenchyma [28] and were thus likely imprinted by their hypoxic *in vivo* microenvironment. Whether these transcriptomic differences result in altered cellular functions remains to be determined in future studies.

BC in the healthy airways are a heterogeneous cell population, including canonical, proliferating and secretory-primed BC subsets [27]. Both our cultured alveolar and airway BC included cells with close transcriptomic similarity to these BC subsets, secretory-primed BC likely arising through spontaneous BC differentiation in culture, following a clear and previously described differentiation gradient [17]. Interestingly, we observed differences in relative abundance between alveolar and airway BC along this differentiation path, leading to a lower amount of secretory-primed alveolar compared to airway BC. Along the differentiation path, a subpopulation showing enrichment for several metallothioneins (MTs), FOS, AGR2 or KRT15 was enriched in alveolar BC. This may indicate a slightly altered differentiation path in alveolar BC, resulting in less secretory-primed BC and could be in line with previous studies, showing disturbed cell differentiation with an accumulation of disease-specific intermediate cell types such as the aberrant basaloid cells in ILD lungs [1, 2]. Interestingly, alveolar BC showed a trend towards reduced expression of BC markers and keratinization genes and increased expression of EMT genes and mesenchymal markers, although to a lesser extent than the aberrant basaloid cells [1, 2]. Furthermore, the genes up-regulated in the altered differentiation path were shown to be associated with cancer (MTs [33], FOS [34], AGR2 [35] or KRT15 [36]), which potentially indicates a contribution of ectopic BC to developing lung carcinoma [37], a risk that is increased in IPF [38]. However, these conclusions are speculative and would need rigorous assessment in dedicated studies. Differences in BC subtype composition between alveolar and airway BC populations may also account for the reduced organoid formation capacity of the alveolar BC population, which possibly contains fewer cells with organoid-forming potential.

Ectopic BC lining HC in the IPF parenchyma appear to preferentially differentiate to MUC5B rather than MUC5AC-expressing cells [39]. Here, we demonstrated that both alveolar and airway BC differentiate to MUC5B and MUC5AC-expressing cells when cultured under identical conditions. However interestingly, cultured alveolar BC maintained a significantly higher tendency to differentiate to MUC5B-expressing cells in organoids. This suggests that both the cellular microenvironment and intrinsic cell-specific properties determine BC fate decision.

Human fibrotic BC instilled into mice have previously been shown to aggravate bleomycin-induced fibrosis [13] and to form HC-like structures within the mouse parenchyma [13, 16]. In this study, we confirmed this trend towards increased fibrosis after instillation of BC in the bleomycin-challenged mouse lung as well as their differentiation to secretory epithelial cells. However, we did not observe differences in the mediated effects of BC from the two origins. Of note, the small sample size of this experiment may limit the robustness of the results and needs to be increased in future studies.

Our dataset was not conclusive regarding alveolar BC origin. Human AT2 cells trans-differentiate to BC through intermediate cells, co-expressing varying levels of AT2 cell- and BC marker genes *in vitro* [7]. To determine, if our cultured alveolar BC could originate from AT2 cells, we compared BC and AT2 cell marker gene expression in alveolar versus airway BC. scRNA-seq analysis revealed an overall trend, although not significant, towards reduced BC marker gene expression in alveolar BC, which was, however, not accompanied with expression of specific AT2 cell marker genes. This observation tends to favour the hypothesis that the source of our cultured alveolar BC would be migrating airway BC. However, it is also possible that alveolar BC could have fully lost AT2 cell traits after trans-differentiation in the lung, or later after expanding them in culture. Furthermore, it is also possible that our cell culture method introduce a selection bias by favouring the outgrowth of cells closely related to classic BC, but not that of intermediate cell types.

## Acknowledgments

The authors acknowledge the microscopy core facility at the Department of Biomedicine in Basel for their support and assistance in ICC/IF, IHC/IF image acquisition and the sciCORE (http://scicore.unibas.ch/) scientific computing center at the University of Basel for performing calculations.

## Author Contributions

**SB**: Cell culture of human primary lung cells, ALI and organoid processing, ICC/IF, IHC/IF, image acquisition and image quantification in cultured cells, human IPF- and mouse tissue slides, immunoblotting, PCR, proliferation assay, scratch assay, data analysis and interpretation; **PK:** Conception and design of the study, cell culture of human primary lung cells, PCR, data analysis and interpretation, writing the manuscript; **JR:** Data analysis and interpretation of scRNA-seq data, revision and editing of the manuscript for important intellectual content; **NA:** bleomycin-mouse model, data analysis and interpretation, revision and editing of the manuscript for important intellectual content; **LP:** bleomycin-mouse model, data analysis and interpretation, revision and editing of the manuscript for important intellectual content; **AP:** bleomycin-mouse model, data analysis and interpretation, revision and editing of the manuscript for important intellectual content; **KEH:** Conception and design of the study, data analysis and interpretation, revision and editing of the manuscript for important intellectual content.

## Data availability

The processed scRNA-seq dataset (the accession is GSE270495) can be viewed on the gene expression omnibus database (GEO).

## Figure legends

**Supplement figure 1:** tSNE showing the distribution of alveolar and airway BC from donor 1 (airway BC-green; alveolar BC-grey), donor 2 (airway BC-dark green; alveolar BC-light blue) and donor 3 (airway BC-lilac; alveolar BC-purple) (a). Percentage of scRNA-seq reads originating from mitochondrial genes (b). Number of detected genes across cells (number of genes with at least one UMI count, on a log-scale) (c).

**Supplement figure 2:** tSNE showing the best matching cell type for each cell from the *SingleR* analysis using as reference the scRNA-seq datasets from Adams et al. [1], Habermann et al. [2], Reyfmann et al. [20], Travaglini et al. [23], Kathiriya et al. [7], Sikkema et al. [25], Murthy et al. [21], Strunz et al. [24] or Deprez et al. [22]. Cell types represented by no more than 5 cells in our dataset were pooled into the category “Others”.

**Supplement figure 3:** Heatmap of the relative expression across clusters, donors, and sample location of selected marker genes.

**Supplement figure 4:** Volcano plots showing the differential expression analysis results between alveolar and airway BC in cluster 1 (a), cluster 3 (b), cluster 4 (c), and cluster 7 (d). Genes colored (beige=decreased, and blue=increased in alveolar BC relative to airway BC) are significant at an FDR threshold of 1%.

**Supplement figure 5:** Functional enrichment analysis amongst differentially expressed genes in cluster 1 (A), cluster 3 (B), cluster 4 (C), and cluster 7 (D) using MSigDB Hallmark gene sets.

**Supplement figure 6:** Functional enrichment analysis amongst differentially expressed genes in clusters 1-4, cluster 6 and cluster 7 using the MSigDB Reactome gene sets from the C2 collection.

## References

1. Adams TS, Schupp JC, Poli S, et al. Single-cell RNA-seq reveals ectopic and aberrant lung-resident cell populations in idiopathic pulmonary fibrosis. Science Advances. 2020;6(28):eaba1983.

2. Habermann AC, Gutierrez AJ, Bui LT, et al. Single-cell RNA sequencing reveals profibrotic roles of distinct epithelial and mesenchymal lineages in pulmonary fibrosis. Science Advances. 2020;6(28):eaba1972.

3. Prasse A, Binder H, Schupp JC, et al. BAL Cell Gene Expression Is Indicative of Outcome and Airway Basal Cell Involvement in Idiopathic Pulmonary Fibrosis. Am J Respir Crit Care Med. 2019;199(5):622–630.

4. Barkauskas CE, Cronce MJ, Rackley CR, et al. Type 2 alveolar cells are stem cells in adult lung. J Clin Invest. 2013;123(7):3025–3036.

5. Rock JR, Onaitis MW, Rawlins EL, et al. Basal cells as stem cells of the mouse trachea and human airway epithelium. Proc Natl Acad Sci U S A. 2009;106(31):12771–12775.

6. Basil MC, Morrisey EE. Lung regeneration: a tale of mice and men. Semin Cell Dev Biol. 2020;100:88–100.

7. Kathiriya JJ, Wang C, Zhou M, et al. Human alveolar type 2 epithelium transdifferentiates into metaplastic KRT5(+) basal cells. Nat Cell Biol. 2022;24(1):10–23.

8. Cai XT, Jia M, Heigl T, et al. IL-4-induced SOX9 confers lineage plasticity to aged adult lung stem cells. Cell Reports. 2024;43(8).

9. Kumar PA, Hu Y, Yamamoto Y, et al. Distal airway stem cells yield alveoli in vitro and during lung regeneration following H1N1 influenza infection. Cell. 2011;147(3):525–538.

10. Yuan T, Volckaert T, Redente EF, et al. FGF10-FGFR2B Signaling Generates Basal Cells and Drives Alveolar Epithelial Regeneration by Bronchial Epithelial Stem Cells after Lung Injury. Stem Cell Reports. 2019;12(5):1041–1055.

11. Kanegai CM, Xi Y, Donne ML, et al. Persistent Pathology in Influenza-Infected Mouse Lungs. Am J Respir Cell Mol Biol. 2016;55(4):613–615.

12. Ray S, Chiba N, Yao C, et al. Rare SOX2(+) Airway Progenitor Cells Generate KRT5(+) Cells that Repopulate Damaged Alveolar Parenchyma following Influenza Virus Infection. Stem Cell Reports. 2016;7(5):817–825.

13. Jaeger B, Schupp JC, Plappert L, et al. Airway basal cells show a dedifferentiated KRT17(high)Phenotype and promote fibrosis in idiopathic pulmonary fibrosis. Nat Commun. 2022;13(1):5637.

14. Hostettler KE, Gazdhar A, Khan P, et al. Multipotent mesenchymal stem cells in lung fibrosis. PLoS One. 2017;12(8):e0181946.

15. Khan P, Fytianos K, Blumer S, et al. Basal-Like Cell-Conditioned Medium Exerts Anti-Fibrotic Effects In Vitro and In Vivo. Frontiers in Bioengineering and Biotechnology. 2022;10.

16. Blumer S, Khan P, Artysh N, et al. The use of cultured human alveolar basal cells to mimic honeycomb formation in idiopathic pulmonary fibrosis. Respir Res. 2024;25(1):26.

17. Khan P, Roux J, Blumer S, et al. Alveolar Basal Cells Differentiate towards Secretory Epithelial- and Aberrant Basaloid-like Cells In Vitro. Cells. 2022;11(11).

18. Ashcroft T, Simpson JM, Timbrell V. Simple method of estimating severity of pulmonary fibrosis on a numerical scale. J Clin Pathol. 1988;41(4):467–470.

19. Carraro G, Langerman J, Sabri S, et al. Transcriptional analysis of cystic fibrosis airways at single-cell resolution reveals altered epithelial cell states and composition. Nat Med. 2021;27(5):806–814.

20. Reyfman PA, Walter JM, Joshi N, et al. Single-Cell Transcriptomic Analysis of Human Lung Provides Insights into the Pathobiology of Pulmonary Fibrosis. Am J Respir Crit Care Med. 2019;199(12):1517–1536.

21. Kadur Lakshminarasimha Murthy P, Sontake V, Tata A, et al. Human distal lung maps and lineage hierarchies reveal a bipotent progenitor. Nature. 2022;604(7904):111–119.

22. Deprez M, Zaragosi LE, Truchi M, et al. A Single-Cell Atlas of the Human Healthy Airways. Am J Respir Crit Care Med. 2020;202(12):1636–1645.

23. Travaglini KJ, Nabhan AN, Penland L, et al. A molecular cell atlas of the human lung from single-cell RNA sequencing. Nature. 2020;587(7835):619–625.

24. Strunz M, Simon LM, Ansari M, et al. Alveolar regeneration through a Krt8+ transitional stem cell state that persists in human lung fibrosis. Nature Communications. 2020;11(1):3559.

25. Sikkema L, Ramírez-Suástegui C, Strobl DC, et al. An integrated cell atlas of the lung in health and disease. Nat Med. 2023;29(6):1563–1577.

26. Hackett NR, Shaykhiev R, Walters MS, et al. The human airway epithelial basal cell transcriptome. PLoS One. 2011;6(5):e18378.

27. Carraro G, Mulay A, Yao C, et al. Single-Cell Reconstruction of Human Basal Cell Diversity in Normal and Idiopathic Pulmonary Fibrosis Lungs. Am J Respir Crit Care Med. 2020;202(11):1540–1550.

28. Xi Y, Kim T, Brumwell AN, et al. Local lung hypoxia determines epithelial fate decisions during alveolar regeneration. Nat Cell Biol. 2017;19(8):904–914.

29. Bodempudi V, Hergert P, Smith K, et al. miR-210 promotes IPF fibroblast proliferation in response to hypoxia. Am J Physiol Lung Cell Mol Physiol. 2014;307(4):L283–294.

30. Tzouvelekis A, Harokopos V, Paparountas T, et al. Comparative expression profiling in pulmonary fibrosis suggests a role of hypoxia-inducible factor-1alpha in disease pathogenesis. Am J Respir Crit Care Med. 2007;176(11):1108–1119.

31. Kierans SJ, Taylor CT. Regulation of glycolysis by the hypoxia-inducible factor (HIF): implications for cellular physiology. J Physiol. 2021;599(1):23–37.

32. Zeng M, Kikuchi H, Pino MS, et al. Hypoxia activates the K-ras proto-oncogene to stimulate angiogenesis and inhibit apoptosis in colon cancer cells. PLoS One. 2010;5(6):e10966.

33. Si M, Lang J. The roles of metallothioneins in carcinogenesis. J Hematol Oncol. 2018;11(1):107.

34. Kuonen F, Li NY, Haensel D, et al. c-FOS drives reversible basal to squamous cell carcinoma transition. Cell Rep. 2021;37(1):109774.

35. Moidu NA, NS AR, Syafruddin SE, et al. Secretion of pro-oncogenic AGR2 protein in cancer. Heliyon. 2020;6(9):e05000.

36. Chen W, Miao C. KRT15 promotes colorectal cancer cell migration and invasion through β-catenin/MMP-7 signaling pathway. Med Oncol. 2022;39(5):68.

37. Königshoff M. Lung cancer in pulmonary fibrosis: tales of epithelial cell plasticity. Respiration. 2011;81(5):353–358.

38. Stella GM, D’Agnano V, Piloni D, et al. The oncogenic landscape of the idiopathic pulmonary fibrosis: a narrative review. Transl Lung Cancer Res. 2022;11(3):472–496.

39. Plantier L, Crestani B, Wert SE, et al. Ectopic respiratory epithelial cell differentiation in bronchiolised distal airspaces in idiopathic pulmonary fibrosis. Thorax. 2011;66(8):651–657.

