## Supplementary figures and images for "Characteristics of ectopic alveolar basal cells relative to airway basal cells in fibrosis"

### Supplement figure 1

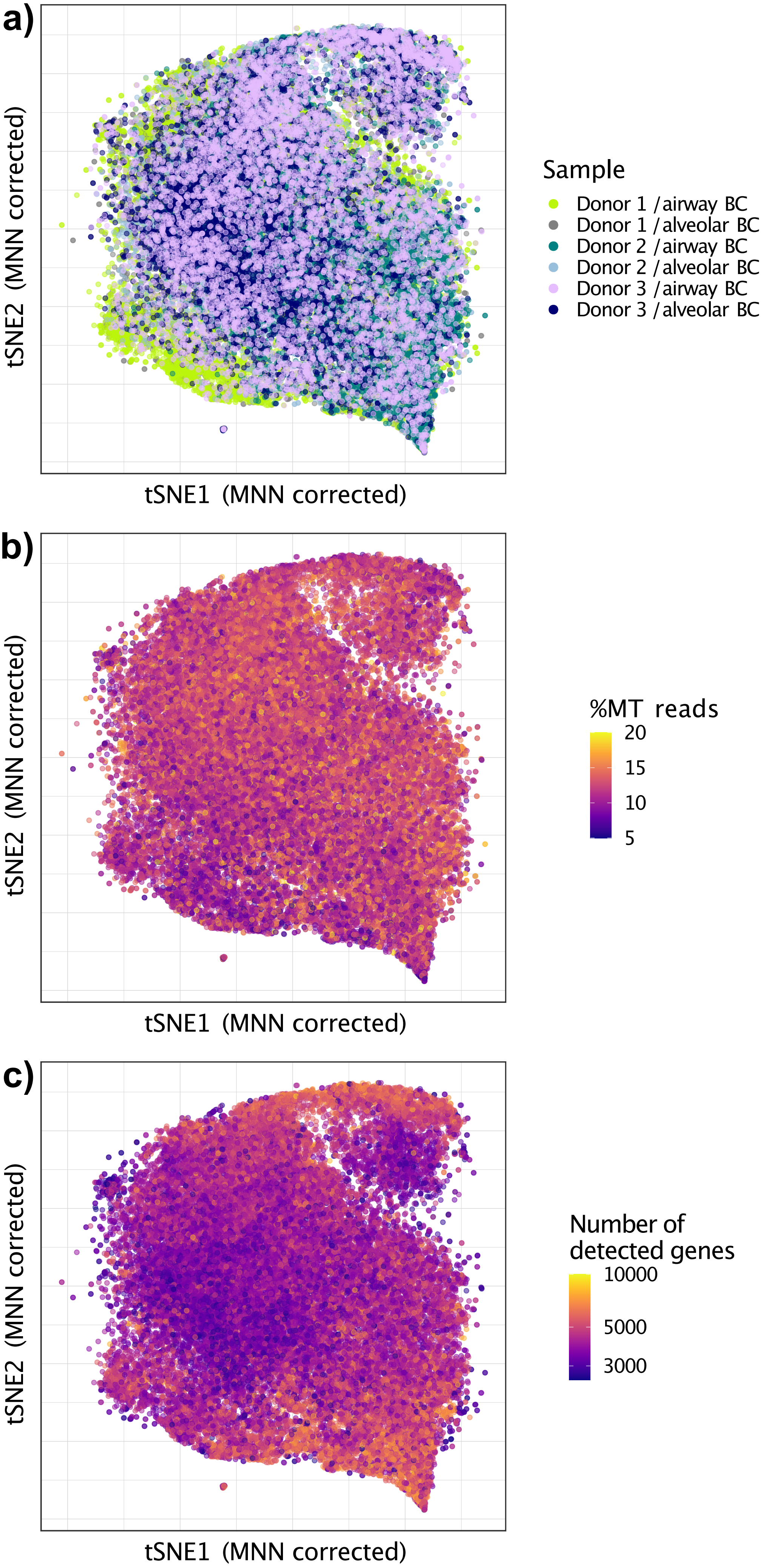

### Supplement figure 2

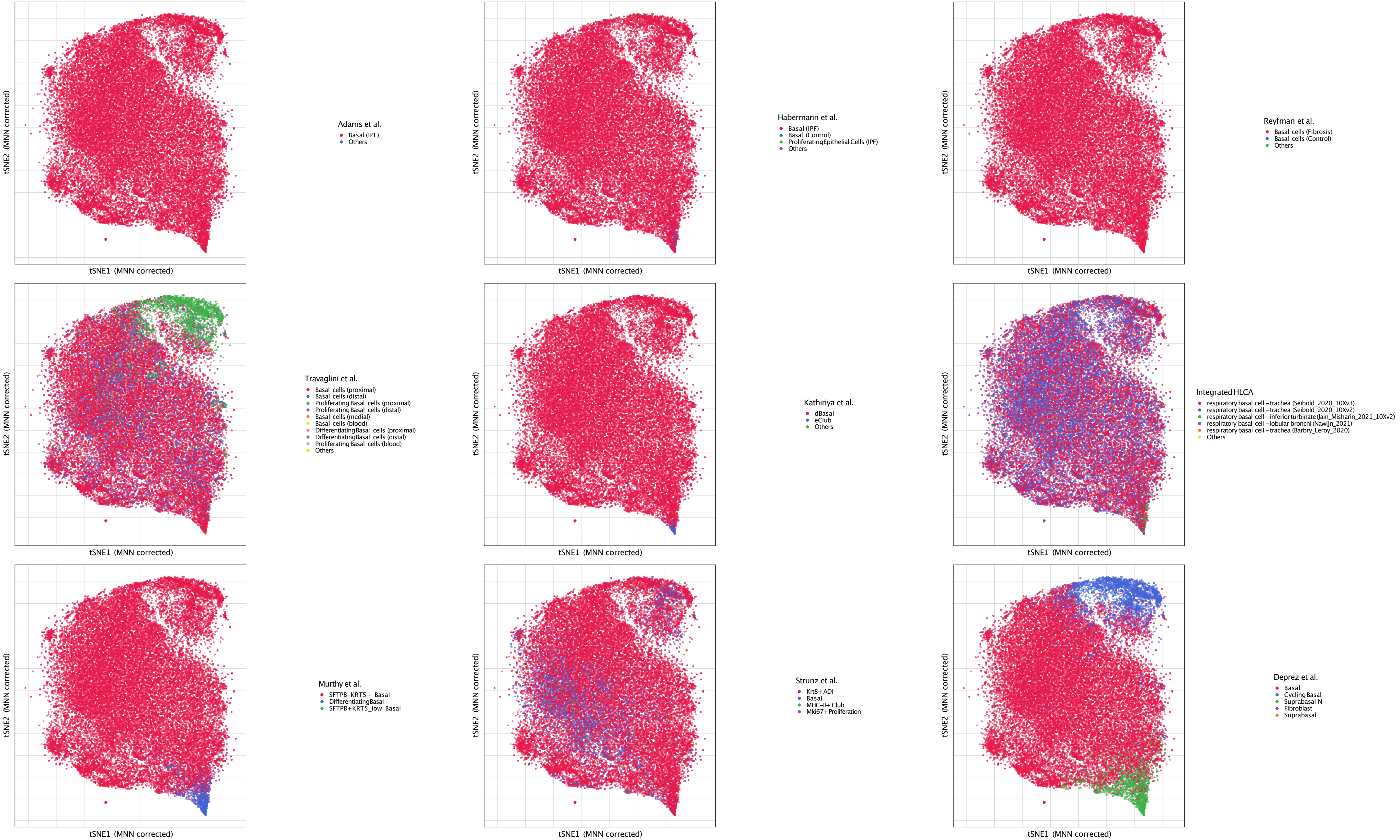

### Supplement figure 3

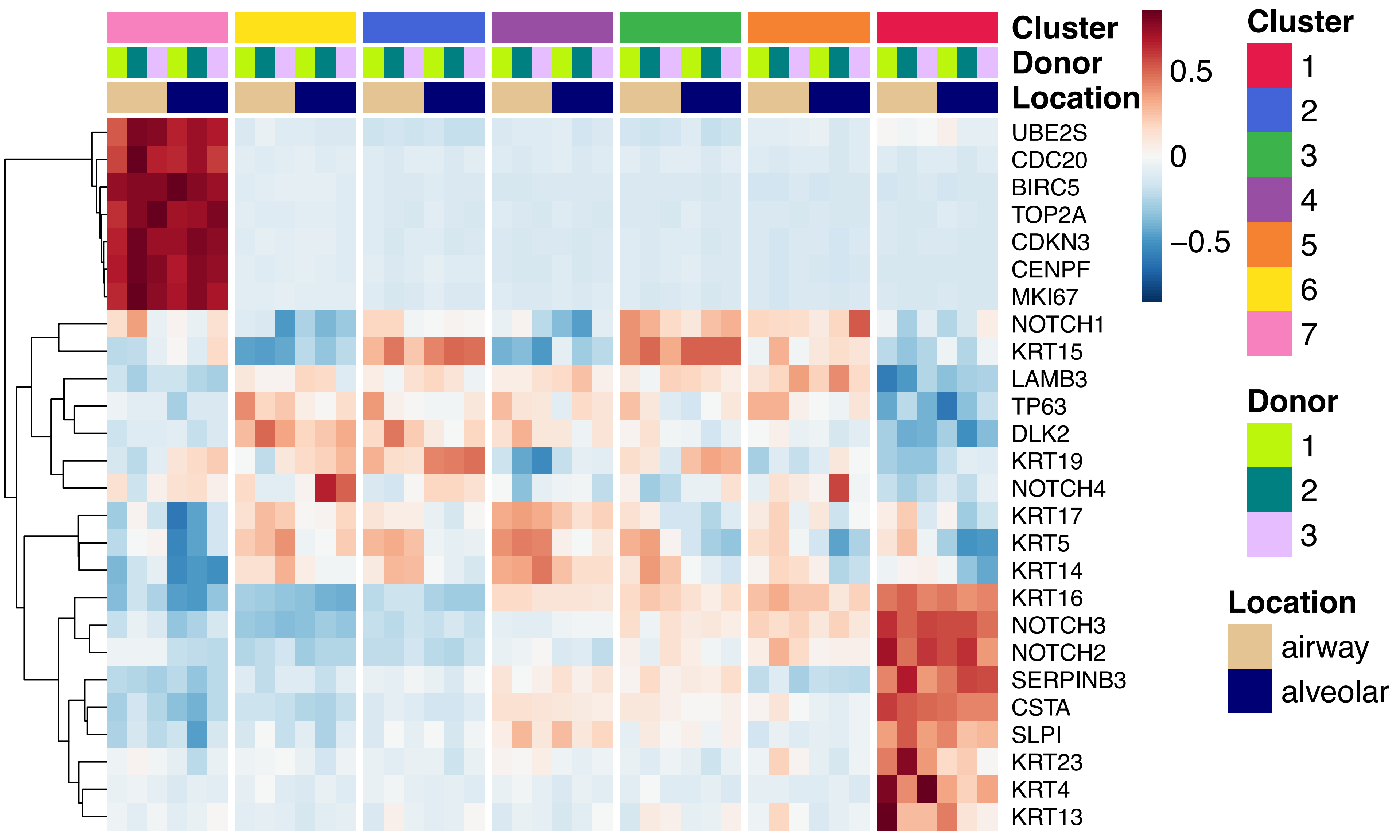

### Supplement figure 4

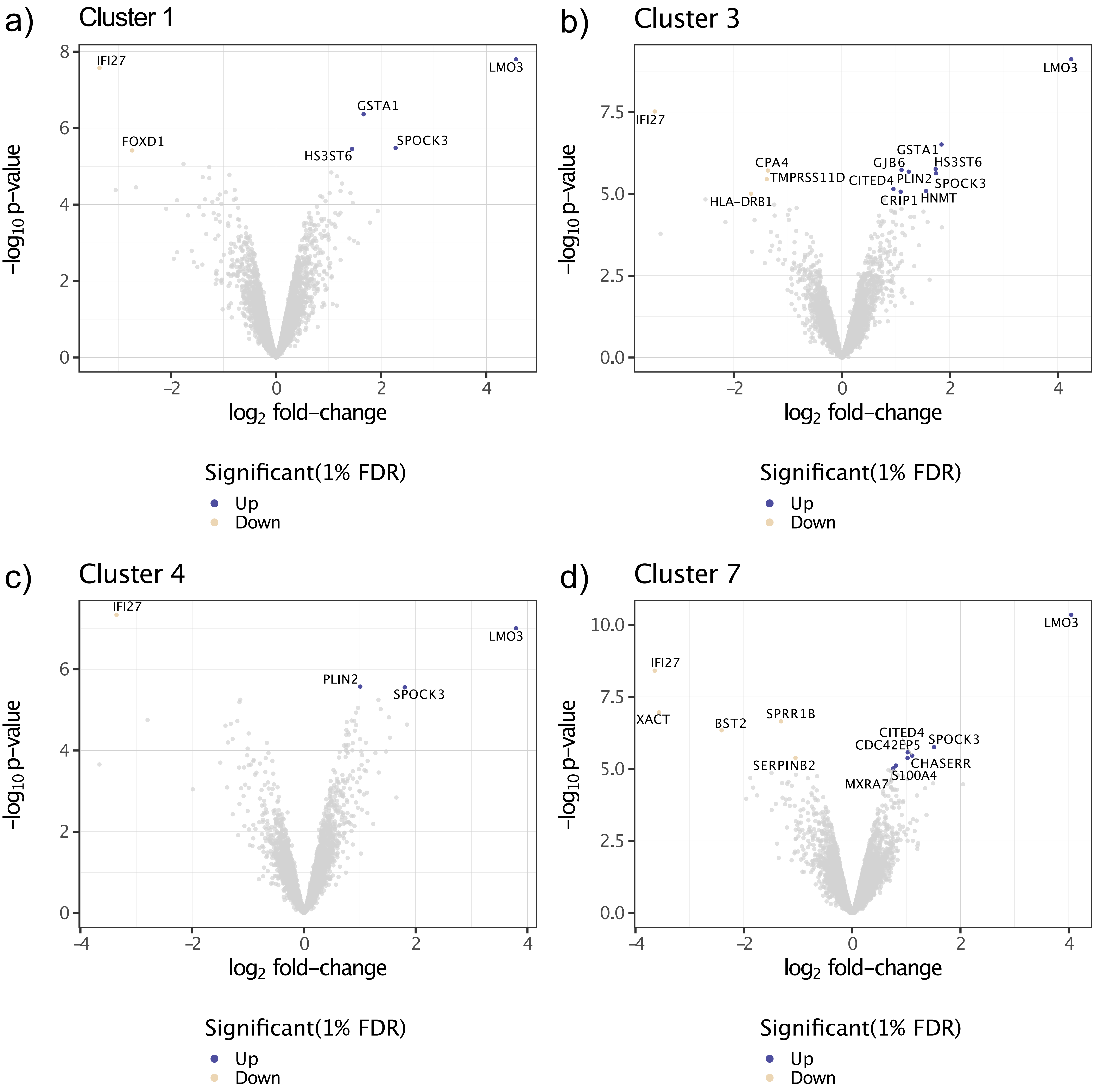

### Supplement figure 5

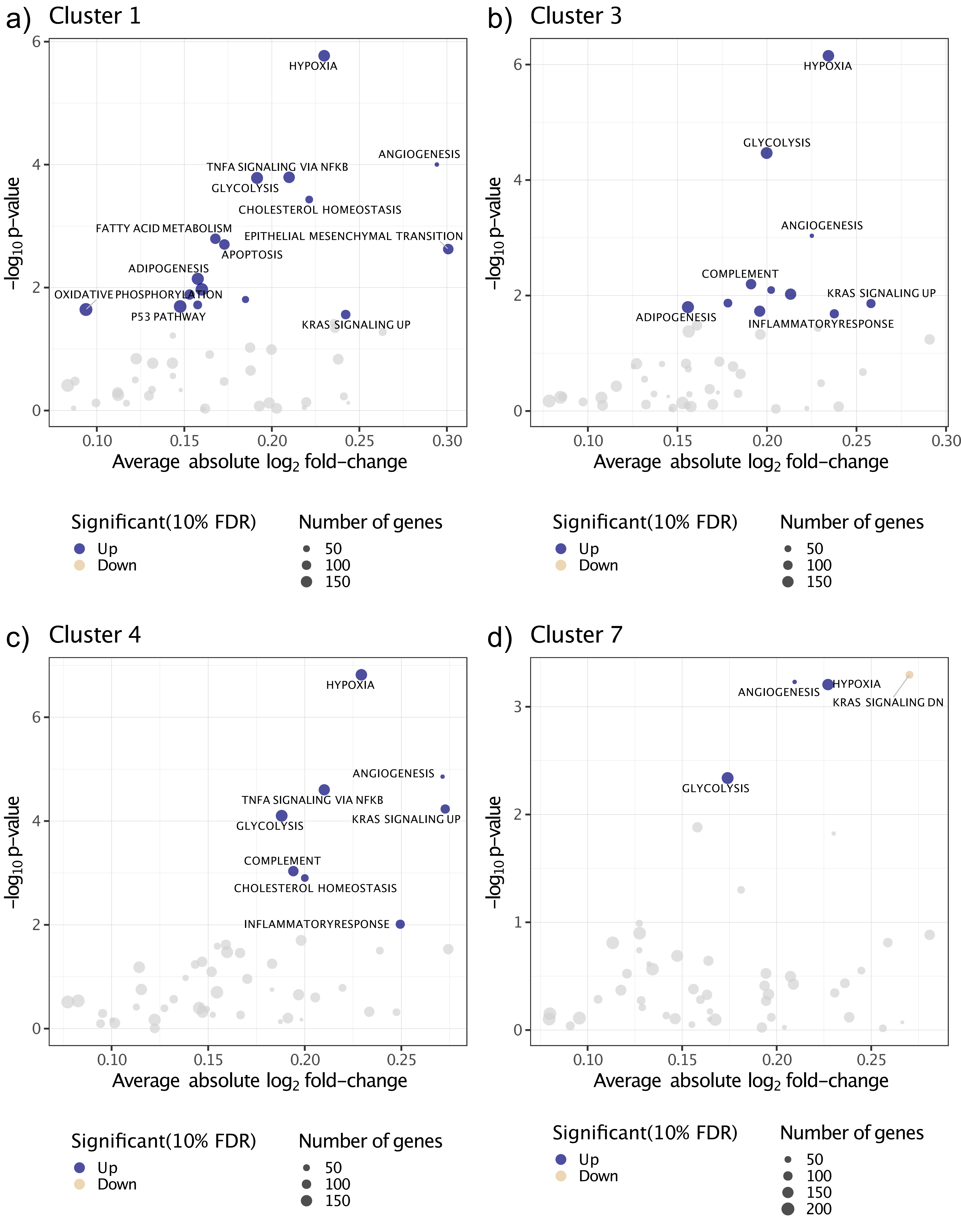

### Supplement figure 6

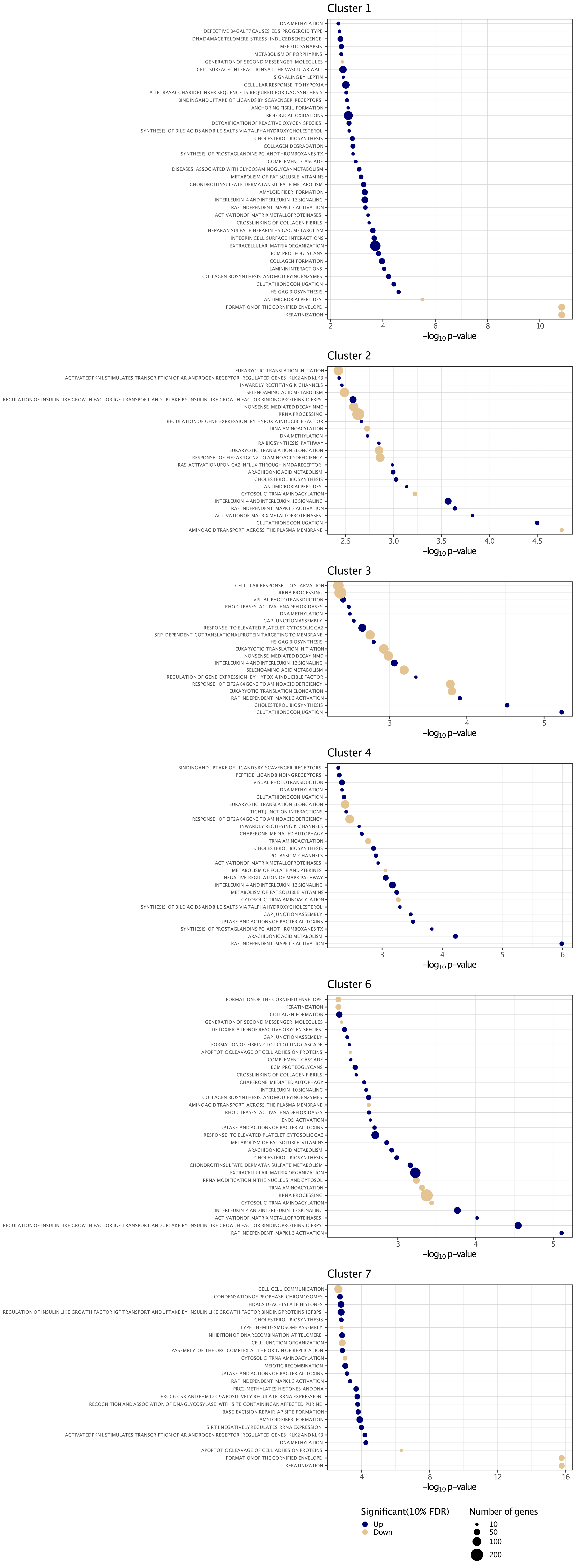
