## Supplement document 1 for "Characteristics of ectopic alveolar basal cells relative to airway basal cells in fibrosis"

*Single-cell RNA-sequencing*

A total of six single cell captures were loaded each onto a channel of a Chromium controller and cDNA and library preparation were performed with a Single Cell 3' v3 Reagent Kit (10X Genomics) according to the manufacturer's instructions. Sequencing was performed on one lane of a flow-cell of the Illumina Novaseq 6000 platform (with 101nt-long R2 reads) at the Genomics Facility Basel of the ETH Zurich, Basel.

The dataset was analyzed by the Bioinformatics Core Facility, Department of Biomedicine, University of Basel. STARsolo (version 2.7.9a) [1] was used to perform sample and cell demultiplexing, and alignment of reads to the human genome (hg38) and UMI counting on gene models from Ensembl 102 (with options "--outFilterType=BySJout --outFilterMultimapNmax=10 --outSAMmultNmax=1 --outFilterScoreMin=30 --soloCBmatchWLtype=1MM_multi_Nbase_pseudocounts --soloUMIfiltering=MultiGeneUMI_CR --soloUMIdedup=1MM_CR --soloType=CB_UMI_Simple –soloStrand=Reverse).

Processing of the UMI counts matrix was performed using the Bioconductor packages DropletUtils (version 1.16.0; function *emptyDrops* using 5000 iterations, the option test.ambient=TRUE, a lower threshold of 100 UMIs and an FDR threshold of 0.1%) [2, 3], scran (version 1.24.1) [4, 5] and scater (version 1.24.0) [6], following mostly the steps illustrated in the OSCA book (http://bioconductor.org/books/3.15/OSCA/) [5, 7]. Cells with less than 5% or more than 20% of UMI counts attributed to mitochondrial genes [8], with less than 10% or more than 30% of UMI counts attributed to ribosomal protein genes, with less than 4000 UMI counts, or with less than 2500 detected genes were excluded. The presence of doublet cells was investigated with the scDblFinder package (version 1.10.0) [9], and suspicious cells were filtered out (doublet score>0.9). Systematic differences between samples were removed using the fastMNN function (d = 50, k = 50) of the batchelor package (version 1.12.30) [10]. tSNE embeddings used for visualization of cells were calculated using the 500 most variable genes and the PCA coordinates or the batch-corrected matrix of low-dimensional coordinates for cells (using a perplexity of 30). Shared nearest-neighbor graph clustering was performed using the Louvain algorithm for finding community structure (k=20). The package SingleR (version 1.10.0) was used for cell-type annotation of the cells [11] using different public lung scRNA-seq datasets as references [12-17] and with the pseudo-bulk aggregation option enabled (aggr.ref=TRUE”).

Differential expression between alveolar and airway BC, controlling for donor-specific differences, was performed using a pseudo-bulk approach, summing the UMI counts of cells from each cluster (excluding cluster 5) in each sample. The aggregated samples were then treated as bulk RNA-seq samples [18]. The package edgeR (version 3.38.4) [19] was used to perform TMM normalization [20] and to test for differential expression with the quasi-likelihood framework (*glmQLFit*). Genes with a false discovery rate (FDR) lower than 1% were considered differentially expressed. Gene set enrichment analysis was performed with the function camera [21] on gene sets from the Molecular Signature Database (MSigDB; version 7.5.1), notably the Hallmark collection and Reactome subset from the C2 collection [22, 23]. We retained only gene sets containing more than 5 genes, and gene sets with a FDR lower than 10% were considered as significant.

1. Dobin, A., et al., *STAR: ultrafast universal RNA-seq aligner.* Bioinformatics, 2013. **29**(1): p. 15-21.
2. Griffiths, J.A., et al., *Detection and removal of barcode swapping in single-cell RNA-seq data.* Nat Commun, 2018. **9**(1): p. 2667.
3. Lun, A.T.L., et al., *EmptyDrops: distinguishing cells from empty droplets in droplet-based single-cell RNA sequencing data.* Genome Biol, 2019. **20**(1): p. 63.
4. Vallejos, C.A., et al., *Normalizing single-cell RNA sequencing data: challenges and opportunities.* Nat Methods, 2017. **14**(6): p. 565-571.
5. Lun, A.T., D.J. McCarthy, and J.C. Marioni, *A step-by-step workflow for low-level analysis of single-cell RNA-seq data with Bioconductor.* F1000Res, 2016. **5**: p. 2122.
6. McCarthy, D.J., et al., *Scater: pre-processing, quality control, normalization and visualization of single-cell RNA-seq data in R.* Bioinformatics, 2017. **33**(8): p. 1179-1186.
7. Amezquita, R.A., et al., *Orchestrating single-cell analysis with Bioconductor.* Nat Methods, 2020. **17**(2): p. 137-145.
8. Ilicic, T., et al., *Classification of low quality cells from single-cell RNA-seq data.* Genome Biol, 2016. **17**: p. 29.
9. Germain, P.L., et al., *Doublet identification in single-cell sequencing data using scDblFinder.* F1000Res, 2021. **10**: p. 979.
10. Haghverdi, L., et al., *Batch effects in single-cell RNA-sequencing data are corrected by matching mutual nearest neighbors.* Nat Biotechnol, 2018. **36**(5): p. 421-427.

11. Aran, D., et al., *Reference-based analysis of lung single-cell sequencing reveals a transitional profibrotic macrophage.* Nat Immunol, 2019. **20**(2): p. 163-172.
12. Adams, T.S., et al., *Single-cell RNA-seq reveals ectopic and aberrant lung-resident cell populations in idiopathic pulmonary fibrosis.* Science Advances, 2020. **6**(28): p. eaba1983.
13. Habermann, A.C., et al., *Single-cell RNA sequencing reveals profibrotic roles of distinct epithelial and mesenchymal lineages in pulmonary fibrosis.* Science Advances, 2020. **6**(28): p. eaba1972.
14. Reyfman, P.A., et al., *Single-Cell Transcriptomic Analysis of Human Lung Provides Insights into the Pathobiology of Pulmonary Fibrosis.* Am J Respir Crit Care Med, 2019. **199**(12): p. 1517-1536.
15. Kathiriya, J.J., et al., *Human alveolar type 2 epithelium transdifferentiates into metaplastic KRT5(+) basal cells.* Nat Cell Biol, 2021.
16. Sikkema, L., et al., *An integrated cell atlas of the lung in health and disease.* Nat Med, 2023. **29**(6): p. 1563-1577.
17. Travaglini, K.J., et al., *A molecular cell atlas of the human lung from single-cell RNA sequencing.* Nature, 2020. **587**(7835): p. 619-625.
18. Lun, A.T.L. and J.C. Marioni, *Overcoming confounding plate effects in differential expression analyses of single-cell RNA-seq data.* Biostatistics, 2017. **18**(3): p. 451-464.
19. Robinson, M.D., D.J. McCarthy, and G.K. Smyth, *edgeR: a Bioconductor package for differential expression analysis of digital gene expression data.* Bioinformatics, 2010. **26**(1): p. 139-40.
20. Robinson, M.D. and A. Oshlack, *A scaling normalization method for differential expression analysis of RNA-seq data.* Genome Biol, 2010. **11**(3): p. R25.
21. Wu, D. and G.K. Smyth, *Camera: a competitive gene set test accounting for inter-gene correlation.* Nucleic Acids Res, 2012. **40**(17): p. e133.
22. Liberzon, A., et al., *The Molecular Signatures Database (MSigDB) hallmark gene set collection.* Cell Syst, 2015. **1**(6): p. 417-425.
23. Subramanian, A., et al., *Gene set enrichment analysis: a knowledge-based approach for interpreting genome-wide expression profiles.* Proc Natl Acad Sci U S A, 2005. **102**(43): p. 15545-50.
