## Supplement table 1 for "Characteristics of ectopic alveolar basal cells relative to airway basal cells in fibrosis"

**Table 1. Patient Characteristics**

| Patient ID | Sex | Age (years) | Clinical Diagnosis |
|---|---|---|---|
| 01 | Female | 67 | Systemic sclerosis-ILD |
| 02 | Male | 79 | IPF |
| 03 | Female | 80 | HP |
| 04 | Male | 83 | HP |
| 05 | Male | 79 | HP |
| 06 | Female | 67 | HP |
| 07 | Female | 62 | Sjögren-associated ILD |
| 08 | Female | 68 | Rheuma-ILD |
| 09 | Male | 31 | Langhans-cell histiocytosis |
| 10 | Male | 44 | Sarcoidosis |
| 11 | Male | 71 | HP |
| 12 | Male | 81 | IPF |
| 13 | Male | 85 | IPF |
| 14 | Male | 58 | Familial pulmonary fibrosis |
| 15 | Male | 78 | MPO-ANCA-associated fibrosis |
| 16 | Male | 63 | IPF |
| 18 | Male | 57 | Systemsclerosis-associated ILD |
| 17 | Male | 56 | Post-COVID fibrosis |
| 18 | Male | 80 | Sarcoidosis |
| 19 | Female | 55 | HP |
| 20 | Female | 79 | Cancer |
| 21 | Male | 70 | IPF |

| | | | |
|---|---|---|---|
| 22 | Female | 56 | ILD (not yet classified) |
| 23 | Male | 73 | Post-COVID fibrosis |
| 24 | Male | 75 | IPF |
| 25 | Male | 84 | HP |
| 26 | Male | 83 | IPF |

*Abbreviations used:* IPF, idiopathic pulmonary fibrosis; ILD, interstitial lung disease; HP, Hypersensitivity pneumonitis; MPO, Myeloperoxidase; ANCA, antineutrophil cytoplasmic antibodies;
